# Novel rRNA-depletion methods for total RNA sequencing and ribosome profiling developed for avian species

**DOI:** 10.1101/2020.09.19.294595

**Authors:** Hanwen Gu, Yu H. Sun, Xin Zhiguo Li

**Author notes:** Corresponding Author Yu H. Sun, Xin Zhiguo Li.

## Abstract

Deep sequencing of RNAs has greatly aided the study of the transcriptome, enabling comprehensive gene expression profiling and the identification of novel transcripts. While mRNAs are of the greatest interest in gene expression studies as they encode for proteins, mRNAs make up only 3-5% of total RNAs, with the majority comprising ribosomal RNAs (rRNAs). Therefore, applications of deep sequencing to RNA face the challenge of how to efficiently enrich mRNA species prior to library construction. Traditional methods extract mRNAs using oligo-dT primers targeting the poly-A tail on mRNAs; however, this approach is not comprehensive as it does not account for mRNAs lacking the poly-A tail or other lncRNAs that we may be interested in. Alternative methods deplete rRNAs, but such approaches require species-specific probes and the commercially available kits are costly and have only been developed for a limited number of model organisms. Here we describe a quick, cost-effective method for depleting rRNAs using custom-designed oligos. We use chickens as an example species for probe design, and the same approach also apply to mice. Using this protocol, we have not only removed the rRNAs from total RNAs for RNA-seq library construction but also depleted rRNA fragments from ribosome-protected fragments for ribosome profiling. Currently, this is the only rRNA depletion-based method for avian species; this method thus provides a valuable resource for both the scientific community and the poultry industry.

## INTRODUCTION

The high-throughput sequencing of cellular RNAs (RNA-seq) or ribosome protected fragments (Ribo-seq) has greatly aided in the characterization of the transcriptome, enabling gene expression studies and the identification of novel transcripts and novel open reading frames [Morlan,et al. (2012), Adiconis, et al. (2013)] {Ingolia et al., 2009, #90822}. Protein coding mRNAs are of the greatest interest in gene expression studies. However, efficient RNA deep sequencing faces significant challenges on how to enrich mRNAs in the sample library, as mRNAs only account for 3-5% of total RNA [Alberts, et al. (2002)]. Therefore, applications of high-throughput sequencing approach to RNAs require methods for enriching mRNAs.

One widely-used method for mRNA enrichment uses of oligo dTs to capture the mRNAs with poly-A tails. However, this approach is incapable of capturing mRNAs that are missing poly-A tails due to degradation, or mRNAs such as histones that use alternative mechanisms for 3’ end processing [López, et al. (2008)]. An alternative method involves the depletion of ribosomal RNAs (rRNAs). Because ribosomes are required for protein synthesis in the cytoplasm and mitochondria, rRNAs are abundant, accounting for ∼80% of total RNA in eukaryotes [Lodish, et al. (2000)]. Cytosolic rRNAs, including 18S, 28S, 5.8S and 5S rRNAs, are processed from 2 rRNA precursors, and the general scheme of their processing is conserved among mammals, invertebrates, yeast, and plants [Tomecki, et al. (2017), Long, et al. (1980)]. One of the rRNA precursors, the 45S rRNA precursor, is transcribed by RNA polymerase I, and then further processed into 18S, 5.8S, and 28S [Winnebeck, et al. (2010)]. The second rRNA precursor, the 5S precursor, is transcribed by RNA polymerase III [Winnebeck, et al. (2010)]. Finally, all mature rRNAs are assembled with ribosomal proteins to form functional small and large subunits. Distinct from cytosolic rRNAs, mitochondria rRNAs, 12S and 16S rRNAs, are encoded by the mitochondrial genome [Rorbach, et al. (2012)], transcribed and processed within mitochondria, and further assembled with ribosome subunits. These mitochondrial rRNAs are responsible for ribosome functions inside mitochondria [Yang, et al. (2014)].

The depletion of rRNAs is not only a preferred mRNA enrichment method for RNA-seq, but also the only option for Ribo-seq. Ribo-seq is a technique to capture ribosome protected fragments (RPFs) on a genome-wide scale, with a sub-codon resolution [Ingolia, et al. (2009)]. RPFs are acquired by treating tissue/cell lysate with an optimized amount of RNase, which degrades the RNAs that are not protected by ribosomes. After running a sucrose gradient, the monosome fraction will be collected to enrich authentic RPFs. Since rRNA is a significant component of ribosomes, the purified RNAs from sucrose gradient also have an abundance of rRNA. Thus, it is also necessary to deplete rRNA before RPF library construction. rRNA-depletion approaches typically use oligos complementary to rRNAs to either extract them or degrade them from the total RNA sample. A major limitation of such antisense oligo-based approaches, however, is that they require species-specific probes. Thus, the commercially available kits are not only costly and but also have been limited to a small number of model organisms, not including chickens.

The chicken has been widely-used for studying various biological questions. Chickens have a sex-determination system different from that of most mammals, wherein males are the homogametic sex (ZZ) and females are the heterogametic sex (ZW). Chicken primordial germ cells (cPGCs) are currently the only type of PGCs that can be cultured *in vitro [*Park, T. S., & Han, J. Y. (2012)], providing a unique opportunity for extensive biochemical studies, including genetic manipulation [Naito, et al (2015), Han, J. Y., & Lee, B. R. (2017)]. Studying chickens benefits not only basic research but also the poultry industry. Chicken is one of the most significant food sources in the world, with over 50 billion chickens being consumed every year [Boyd, et al. (2013)]. Furthermore, given the adaptive radiation of birds (over 10,000 species), chickens are valuable models for endangered avian species conservation [Muir, et al. (2003)]. Investigating chicken genomes and transcriptomes may, therefore, shed light on avian evolutionary processes and contribute to conservation biology [Li et al. (2019)].

Here we focus on developing a method for depleting rRNAs from total RNA using oligos specially designed for chickens. These oligos are antisense to cytosolic and mitochondrial rRNAs, and the resulting DNA-RNA hybrids can be removed using RNase H. RNase H is an endoribonuclease that recognizes DNA-RNA hybrid molecules and hydrolyzes the RNA strand [Inoue, et al. (1987)]. DNase is then used to digest the remaining DNA in the mixture. We have succeeded in using this protocol not only to remove rRNAs from total RNAs for RNA-seq library construction, but also to remove rRNA fragments for Ribo-seq.

Every step of the protocol has been optimized. Specifically, we optimized the ratio between RNAs and oligo probe, the ideal brand of RNase H enzyme, the amount of RNase H, as well as the treatment time and temperature. For total RNA, we monitored our depletion efficiency by the ratio of GAPDH to rRNAs and the success of depletion is shown by bioanalyzer profiles. For RPFs, we adopted the optimized condition from total RNA rRNA depletion, but further optimized the temperature of RNase H treatment as the annealing differs by fragment length. We monitored the depletion efficiency by Ribo-seq and the ratio of rRNA reads in the library. Our strategy has been successfully applied to chickens, and we envision no problems in applying it to other species.

Our protocol is cost-effective and, to the best of our knowledge, is currently the only rRNA depletion-based method for avian species, thus providing a valuable resource for the community. This manuscript outlines the steps for extracting total RNA from animal tissue, depleting rRNAs, and purifying the enriched mRNA sample.

## MATERIALS AND METHODS

The RNA depletion protocol outlined here consists of 2 parts: the oligo-based rRNA depletion (fig 1) and the bioinformatic analysis of the result (https://github.com/LiLabZhaohua/RiboSeqPipeline).

**Figure 1:**
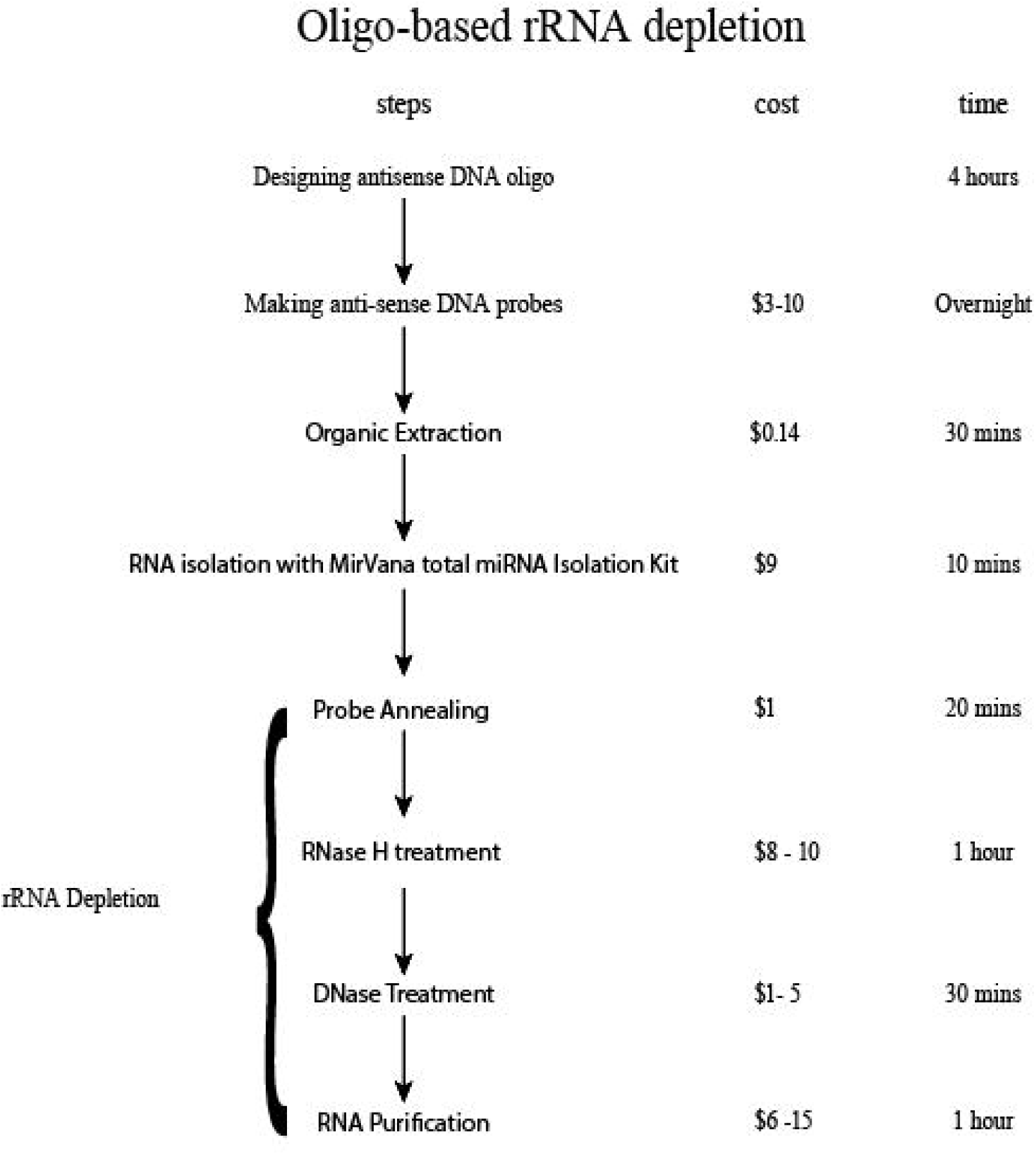
A flowchart summarizing the protocol.

### 1. Designing the antisense oligo

To design the antisense oligo, we downloaded the rRNA sequences of both cytosolic ribosomes and mitochondrial ribosomes in chicken species from NCBI. These are mitochondrial rRNAs including the *16S* rRNA (XR_003076910.1), and *12S* rRNA (MH879470), and cytosolic rRNAs which for eukaryotes include *18S* rRNA (AF173612.1), *28S* rRNA (XR_003078040.1), *5*.*8S* rRNA (XR_003078039.1) and *5S* rRNA (XR_003076910.1). In order to minimize the chances of mismatch, for species not close related to chicken, a specific set of oligos should be designed and synthesized according the following steps.

DNA oligos were designed by converting the original sequences to their reverse complements and splitting the full-length sequences into 50 nt non-overlapping windows using a python script [https://github.com/LiLabZhaohua/rRNADepletion].

1.1. To minimize potential oligo mismatch pairing with other RNAs, the basic local alignment tool (BLAST) was used to find possible pairing with other RNAs in chickens. We confirmed that these oligos do not have significant targets in the chicken transcriptome.
1.2. A pool of DNA probes was made, after receiving the oligos synthesized by Integrated DNA Technologies (Coralville, Iowa 52241), by mixing the *5*.*8S, 28S, 18S, 16S, 12S* and *5S* oligos to a final concentration of 0.5 μM for each oligo.

### 2. Total RNA isolation using mirVana total miRNA isolation kit

This step is done with the mirVana total miRNA isolation kit (Thermo Fisher cat. AM1560).

#### Frozen tissue or extremely hard tissues

2.1. Remove the tissue from the -80 °C refrigerator.
2.2. Cut out 25 mg of frozen tissue and place into a 1.7 uL tube.
2.3. Add 300 uL Lysis/Binding Buffer into a plastic weigh boat or tube on ice.
2.4. Grind frozen tissue into a powder using a homogenizer.
2.5. Add 30 uL miRNA Homogenate Additive to a tube. (Note: It is important to add the Homogenate additive immediately to the tube after the tissue is ground as it prevents the degradation of RNA.)
2.6. Leave mixture on rotator for 10 minutes.

### Organic Extraction

Add 300 uL of Acid Chloroform (Thermo Fisher cat. AM9722).

Withdraw the bottom layer, avoiding the upper phase as this is mostly aqueous buffer.

2.7. Vortex the mixture for 60 seconds to mix well.
2.8. Centrifuge the mixture for 5 minutes at 16,900 RCF at room temperature. (Note: The separation between the top and bottom phase should be clear. If the interphase is not compact, repeat the centrifugation process.)
2.9. Transfer the upper layer to a new 1.7 uL tube. It should be approximately 300 uL. It is very important to not disturb the bottom phase since it is mostly tissue and acid phenol.

#### RNA Isolation Procedure

Wash Solution 1 and Solution 2/3 should be pre-mixed with ethanol according to the manual. At the end of the final stage, RNA can be eluted with either nuclease-free water or in the Elution buffer provided by the kit.

2.10. Add 375 uL of room temperature 100% ethanol to the aqueous phase.
2.11. Place column on a collection tube.
2.12. Pipet the mixture to the column.
2.13. Centrifuge for 30 seconds at 9300 RCF.
2.14. Discard flow through.
2.15. Add 700 uL miRNA Wash Solution 1 (Pre-mixed with ethanol).
2.16. Centrifuge for 10 seconds.
2.17. Discard flow through.
2.18. Add 500 uL miRNA wash Solution 2/3 (Pre-mixed with ethanol).
2.19. Centrifuge for 10 seconds.
2.20. Discard flow through.
2.21. Add 500 uL miRNA wash Solution 2/3 (pre-mixed with ethanol).
2.22. Centrifuge for 10 seconds.
2.23. cDiscard flow through.
2.24. Centrifuge for another 3 minutes to remove the liquid residual.
2.25. Pre-heat the nuclease-free water to 95 °C.
2.26. Transfer the column into a new collection tube.
2.27. Add 100 uL pre-heated Elution Buffer or nuclease-free water.
2.28. Spin for 4 minutes at 11300 RCF at room temperature.
2.29. Collect elutes and store at -80 °C.

### 3. rRNA depletion

For this step, the enrichment for mRNAs and RPFs follows the same steps **except** for the temperature for RNase H treatment and the last step, in which the depletion from total RNAs uses the RNA Clean & Concentrator kit (ZYMO Research cat. R1013) protocol, while the depletion from RPF uses the Acid Phenol protocol.

#### Depletion Preparation

*In this step, we prepare three solutions that will be used for rRNA depletion later*.

3.1 For the probe oligo, prepare the 5X rRNA oligo Hybridization Buffer and 10X RNase H and aliquot for use.
3.2 Prepare 5X rRNA oligo Hybridization Buffer: mix 100 mM Tris-HCl with 200 mM NaCl.
3.3 Prepare 10X RNase H Digestion Buffer: mix 500 mM Tris-HCl with 1 M NaCl and 200 mM MgCl_2_.

### 3.1 rRNA depletion from total RNA

#### Probe annealing

3.1.1 Pipette 2 μg of RNA and 15 μL DNA oligo probe into a tube, adjusting the volume of 5X rRNA oligo Hybridization Buffer and water according to Table II.

**Table I:** Read statistics of Ribo-seq libraries constructed with different annealing temperature This is RPF data, it could be 99%. We chose the optimized temperature.

| Name | Total Reads | rRNA Mapping Reads | % rRNA Mapping Reads |
| --- | --- | --- | --- |
| Sample 1 - 30 | 763,350 | 372,204 | 48.8 |
| Sample 2 - 37 | 165,239 | 63,454 | 38.4 |
| Sample 3 - 42 | 743,014 | 301,481 | 40.6 |
| Sample 4 - 45 | 937,034 | 388,145 | 41.4 |
| Sample 5 - RT | 973,777 | 475,268 | 48.8 |

**Table II:** Probe annealing system
3.1.2. You may adjust the volume and choose one system in Table II, or use a larger total volume, but a total volume over 45 μL is not suggested.
3.1.3 You may also dilute the system, using the 1X rRNA oligo Hybridization Buffer, after you mix all the components (not over 45 μL for 2μg RNA).
3.1.4. Use the PCR machine for annealing: Heat to 95 °C for 2 minutes and slowly cool down (−0.1 °C/sec) to 22 °C, then incubate at 22 °C for additional 5 minutes.

|  |  |  |  |
| --- | --- | --- | --- |
|  | 1 | 2 | 3 |
| RNA | 2 ug |  |  |
| 5X rRNA oligo Hybridization Buffer | 5 uL | 6uL | 7uL |
| DNA oligo probe | 15 uL |  |  |
| ddH2O | to 25 uL | to 30 uL | to 35 uL |

#### RNase H treatment

3.1.5 Turn on the PCR machine and set the temperature to 45 °C.
3.1.6 Add the solutions and Invitrogen RNase H (Thermo Fisher cat. 18021071) listed in Table III, according to the volumes of annealing.

**Table III:** RNase H treatment system
3.1.7 Incubate at 45 °C for 1 hour in the PCR machine.

|  | 1 | 2 | 3 |
| --- | --- | --- | --- |
| 10X RNase H<br>Digestion Buffer | 3 uL | 3.5 uL | 4 uL |
| Invitrogen RNase H | 1 uL |  |  |
| ddH2O | 1 uL | 0.5 uL | 0 uL |
| Total volume | 30 uL | 35 uL | 40 uL |

#### DNase treatment

3.1.8 Take out samples from the PCR machine and adjust the PCR machine to 37 °C.
3.1.9 Immediately add the components listed in Table IV. Note that regardless of which system you chose, the total volume should be 50 uL.

**Table IV:**
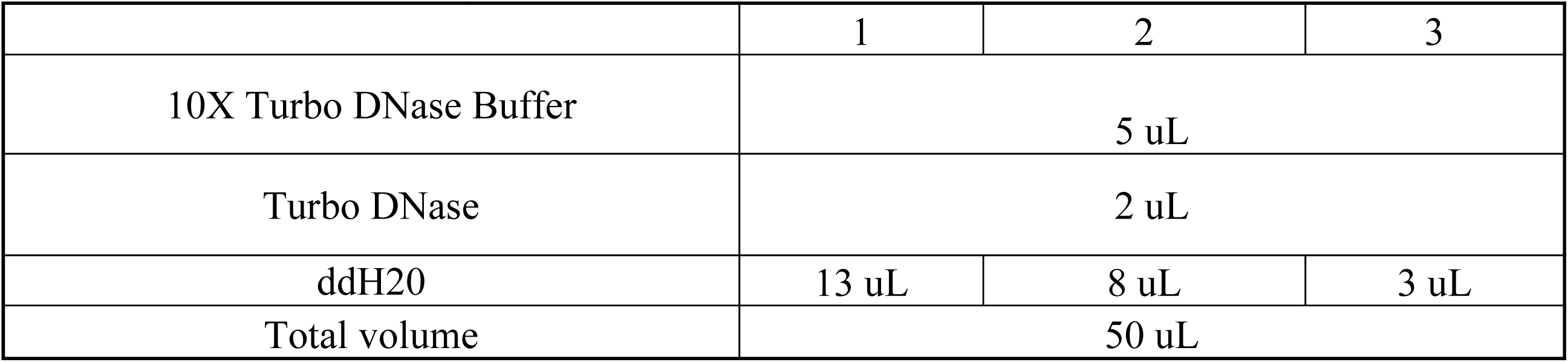
DNase treatment system
3.1.10. Incubate at 37 °C for 30 minutes in the PCR machine.

#### RNA purification

This step uses the RNA Clean & Concentrator kit (Thermo Fisher cat. R1013) protocol. Prepare adjusted RNA Binding buffer by mixing equal volumes of buffer and ethanol.

3.1.11. Transfer 50 uL of RNA to a new 1.7 mL tube.
3.1.12. Add 50 ul water to the tube.
3.1.13. Add 200 uL adjusted binding buffer to the tube.
3.1.14. Transfer the mixture into a column.
3.1.15. Centrifuge for 30 seconds at 16,900 RCF at room temperature.
3.1.16. Discard the flow-through.
3.1.17. Add 200 uL RNA Pre-Wash.
3.1.18. Centrifuge for 30 seconds at 16,900 RCF at room temperature.
3.1.19. Discard the flow-through.
3.1.20. Add 200 uL RNA Wash Buffer.
3.1.21. Centrifuge for 30 seconds at 16,900 RCF at room temperature.
3.1.22. Discard the flow-through.
3.1.23. Add 200 uL RNA Wash Buffer.
3.1.24. Centrifuge for 2 minutes at 16,900 RCF at room temperature.
3.1.25. Discard the flow-through.
3.1.26. Transfer the columns into new 1.5 mL tubes (RNase Free tube).
3.1.27. Add 15 uL DNase/RNase-Free water directly to the column matrix.
3.1.28. Let the columns sit at room temperature for at least 1 minute.
3.1.29. Centrifuge for 30 seconds at 16,900 RCF at room temperature.
3.1.30. Keep the flow through in the tube and store the tube at -80 °C immediately.

### 3.2. rRNA depletion from ribosome protected fragments (RPFs)

#### Probe annealing

3.2.1. Pipette 2 μg of RNA and 15 μL DNA oligo probe into a 1.7 ml tube.
3.2.2. Adjust the volume of 5X rRNA oligo Hybridization Buffer and water according to Table II.
3.2.3. You may adjust the volume and choose one system in Table II, or use a larger total volume, but a total volume over 45 μL is not suggested.
3.2.4. You may also dilute the system, using the 1X rRNA oligo Hybridization Buffer, after you mix all the components (not over 45 μL for 2μg RNA).
3.2.5. Use the PCR machine rRNA program for annealing: Heat to 95 °C for 2 minutes and slowly cool down (−0.1 °C/sec) to 22 °C, then incubate at 22 °C for additional 5 minutes.

#### RNase H treatment

3.2.6. Turn on the PCR machine and set the temperature as 37 °C for RPF for optimal results. * We have compared the treatment temperature at room temperature, 30 °C, 37 °C, 42 °C, and 45 °C, and found that 37 °C has the best removal based on our sequencing results (Table I).
3.2.7. Add the solutions and RNase H (Thermo Fisher cat. 18021071) listed in Table III, according to the volumes of annealing.
3.2.8. Incubate at 37 °C for 1 hour on PCR machine.

#### DNase treatment

3.2.9. Remove the samples from PCR machine and adjust the PCR machine to 37 °C.
3.2.10. Immediately add the components listed in Table IV. (Note: The total volumes should all be 50 uL.)
3.2.11. Incubate at 37 °C for 30 minutes in the PCR machine.

#### Purification of RPF

3.2.12. Transfer 50 uL RNA to a 1.7 mL tube.
3.2.13. Add 200 μL ddH2O.
3.2.14. Add 300 μL Acid Phenol Chloroform (Thermo Fisher cat. AM9722). Note the total volume should be around 550 uL.
3.2.15. Vortex for 1 minute to mix well.
3.2.16. Spin at 20,100 RCF for 15 minutes at room temperature.
3.1.17. Withdraw the upper phase (around 250 uL) without disturbing the lower phase and transfer the solution to a new 1.7 mL tube.
3.2.18. Add 1 μL glycogen (Sigma Aldrich cat. 10901393001).
3.2.19. Add 3 volumes (around 750 uL) of 100% ethanol.
3.2.20. Leave at 4 °C for an hour, or -20 °C overnight. White precipitate should be visible.
3.2.21. Spin down at 16,900 RCF for 30 minutes at 4 °C.
3.2.22. Wash with 200 uL of 70% ethanol.
3.2.23. Centrifuge at 16,900 RCF for 5 minutes at highest speed at 4 °C.
3.2.24. Pour out the liquid.
3.2.25. Centrifuge at 16,900 RCF for another 3 minutes at highest speed at 4 °C.
3.2.26. Pipette out all the liquid and leave only the pellet in the tube.
3.2.27. Dissolve in water at room temperature.
3.2.28. Store at -80 °C.

## Data Analysis

We analyzed the Ribo-seq as previously described [Sun, Y. H., etc., (2020)]. Ribo-seq pipeline for chicken was developed for this study, which is available in the following link (https://github.com/LiLabZhaohua/RiboSeqPipeline).

## RESULTS

In this protocol, we describe a simple method for enriching mRNAs from animal tissue using an rRNA depletion-based approach. Each key step of the protocol reported above has been optimized, including the ratio between RNAs and oligo probes, the ideal brand of RNase H enzyme, the amount of RNase H, as well as the treatment time and temperature. This is, to the best of our knowledge, the first rRNA depletion method developed for avian species. Although the probes are designed to target chicken rRNAs, considering the conservation of rRNA sequences, we expect that our probes can be used for diverse avian species.

For total RNAs, we used the ratio of 28S to GAPDH to measure the depletion efficiency. We first compared efficiency of three commercial RNase H enzymes – RNase H (NEB cat. M0297S) and thermostable RNase H (NEB cat. M0523S) from NEB, and RNase (Thermo Fisher cat. 18021071) from Invitrogen with 1 ul probe, 1 ul RNase H treated for 30 mins at 37 °C and 45 °C. The smallest Relative quality value with respect to the control value for RNase H from Invitrogen indicates the best result at 37 °C. Then we compared the Invitrogen RNase H and NEB RNase H with 1 ul and 2 ul at 37 °C and 45 °C with 3 μl probe for 1 hour of RNase treatment and then 30 mins of DNase treatment. The result indicates that Invitrogen has the best efficiency at 45 °C. After the optimal enzyme chosen, the optimal amount of probe was also investigated with 1μl Invitrogen RNase Treatment at 45 °C for 1 h and then DNase treatment for 30 mins. Different ratios of the RNA and probe, including 1:0, 1:1, 1:2, and 1:3 were tested. In the end, the optimal condition for RNA depletion is 1 μg RNA: 3 μl probe, 0.2 μl Invitrogen RNase H treatment at 45°C for 1 h, and DNase treatment at 37°C for 30 min. The result was expressed as the ratio of 28S to GAPDH, decreased > 4,700 fold from 706 to 0.15 after rRNA depletion. The success of rRNA depletion is also demonstrated by the bioanalyzer results, where the 18S and 28S rRNA peaks disappeared after depletion at the optimal condition which is 1 μg RNA:3 μl probe, with 0.2 μl Invitrogen RNase treatment at 45°C for 1 hour, and then DNase treatment at 37°C for 30 mins. The experiment is considered successful when all ribosomal RNA, including 18S and 28S, are depleted by the end of the procedure. The lack of any rRNA peaks demonstrates the complete removal of rRNA in the rRNA-depletion process (Figure 2B & Figure 2E are two replicates of the samples after depletion). The small peaks that can be seen typically correspond to some small RNA and mRNA.

**Figure 2:**
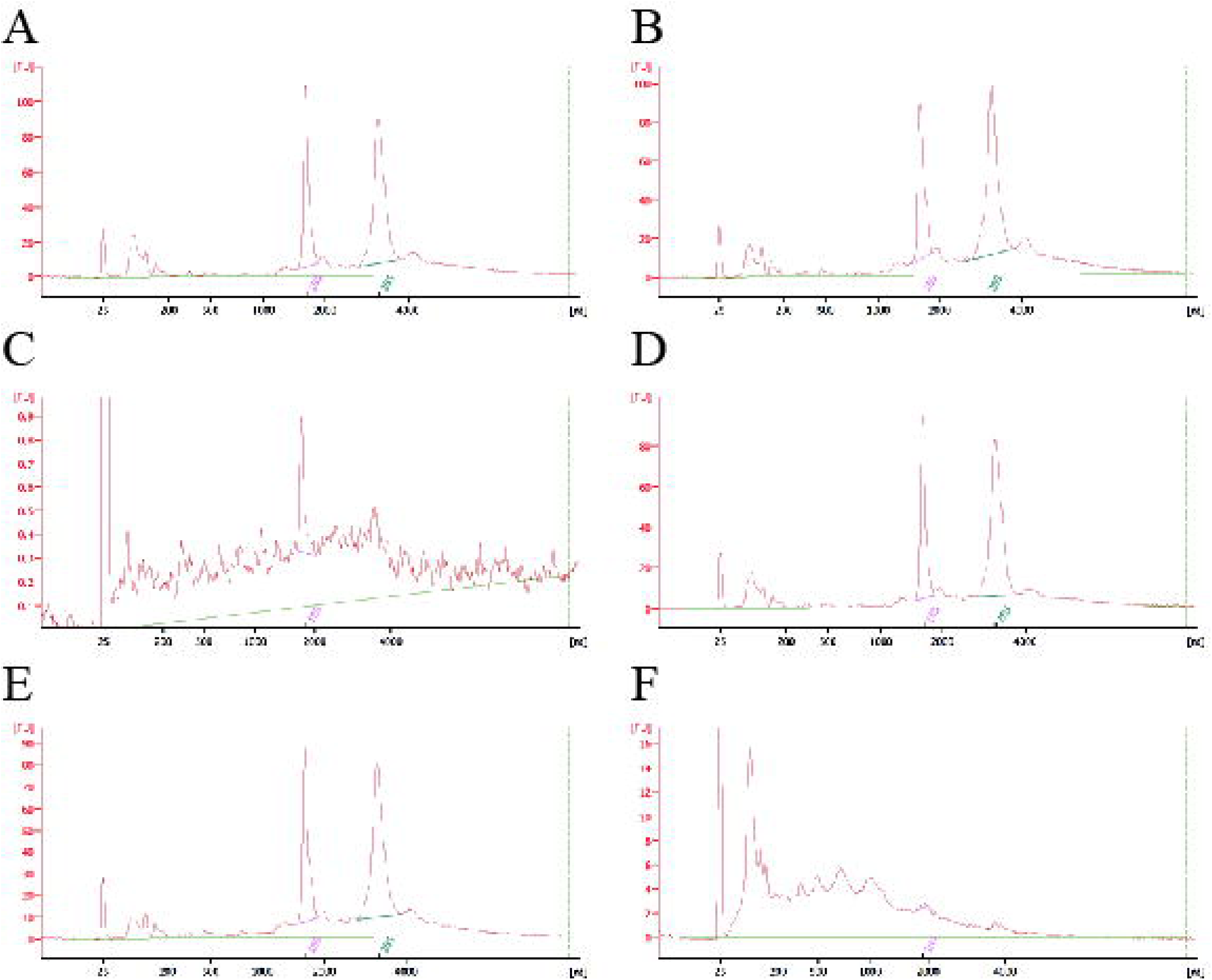
Total RNA bioanalyzer profile, where x axis is RNA size in nucleotides and y axis is the arbitrary fluorescence unit. The profile is typically expressed with 2 major peaks corresponding to the 18S and 28S ribosomal RNA (Figure 2A & Figure 2D). 3 parameters including the RIN number, 260/230 ratio, and 260/280 ratio can be used to measure RNA quality, as well as the effect of DNase treatment. The RNA integrity number (RIN) is an index for assigning integrity values to RNA measurements. No significant change should be observed for a successful experiment before and after DNase treatment. Figure 2A & 2B and Figure 2D & 2E are two sets of replicates that have a stable RIN before and after DNase treatment indicating the success of the treatment. The 260/230 values for “pure” nucleic acid are often used to examine whether contamination such as salt or protein exists in the sample or to measure the purity of nucleic acid before and after. Expected 260/230 values are commonly in the range of 2.0-2.2. If the ratio is appreciably lower, it may indicate the presence of contaminants which absorb at 230 nm. The ratio would be decrease largely due loss of rRNA. Sample A & B and D&E are two sets of replicates that have a stable 260/230 ratio before, and after which are considered successfully treated with DNase. The 260/280 values indicate the purity of RNA. A ratio of ∼2.0 is generally accepted as “pure”, while pure DNA has a ratio close to 1.8. Sample A & B and D&E are two sets of replicates that have a stable 260/280 ratio before and after DNase treatment. This result reveals the successful removal of rDNA oligos. The RNA concentration within the sample is another parameter used to evaluate the success of the RNA depletion. The value should decrease largely after depletion as 90% of the RNA, which is ribosomal RNA, is now depleted. The decrease of the RNA concentration indicates that sample A&C and D&F are two successful treatments.

For Ribo-seq, we used the percentage of rRNA fragment in the sequencing results to measure the depletion efficiency. Because the RNA length differs between full-length mRNAs and RPFs, likely to affect the annealing temperature, we therefore constructed Ribo-seq library with the RNase H treatment temperature at RT, 30°C, 37°C, 42°C, and 45°C respectively. Table I listed the read statistics of the Ribo-seq libraries showing the number of total reads and number of reads mapping to the rRNA sequences. Based on the result in Table I, the optimal result of RPF depletion is seen at 37 °C with the lowest percentage of reads coming from rRNAs. Therefore, 37 °C is used for RPF depletion protocol.

Our results are consistent with what have been reported in mice [Adiconis, et al. (2013)], which indicate the RNase H method is robust and capable of depleting rRNAs from low input samples. As described before [Adiconis, et al. (2013)] that compared to Ribo-Zero, this method has a significantly higher efficiency in rRNA removal and slightly better evenness of coverage, we also found the chicken RNA-seq signals evenly distribute throughout the mRNAs as shown by the metagene analysis of a chicken RNA-seq results (Figure 3).

**Figure 3:**
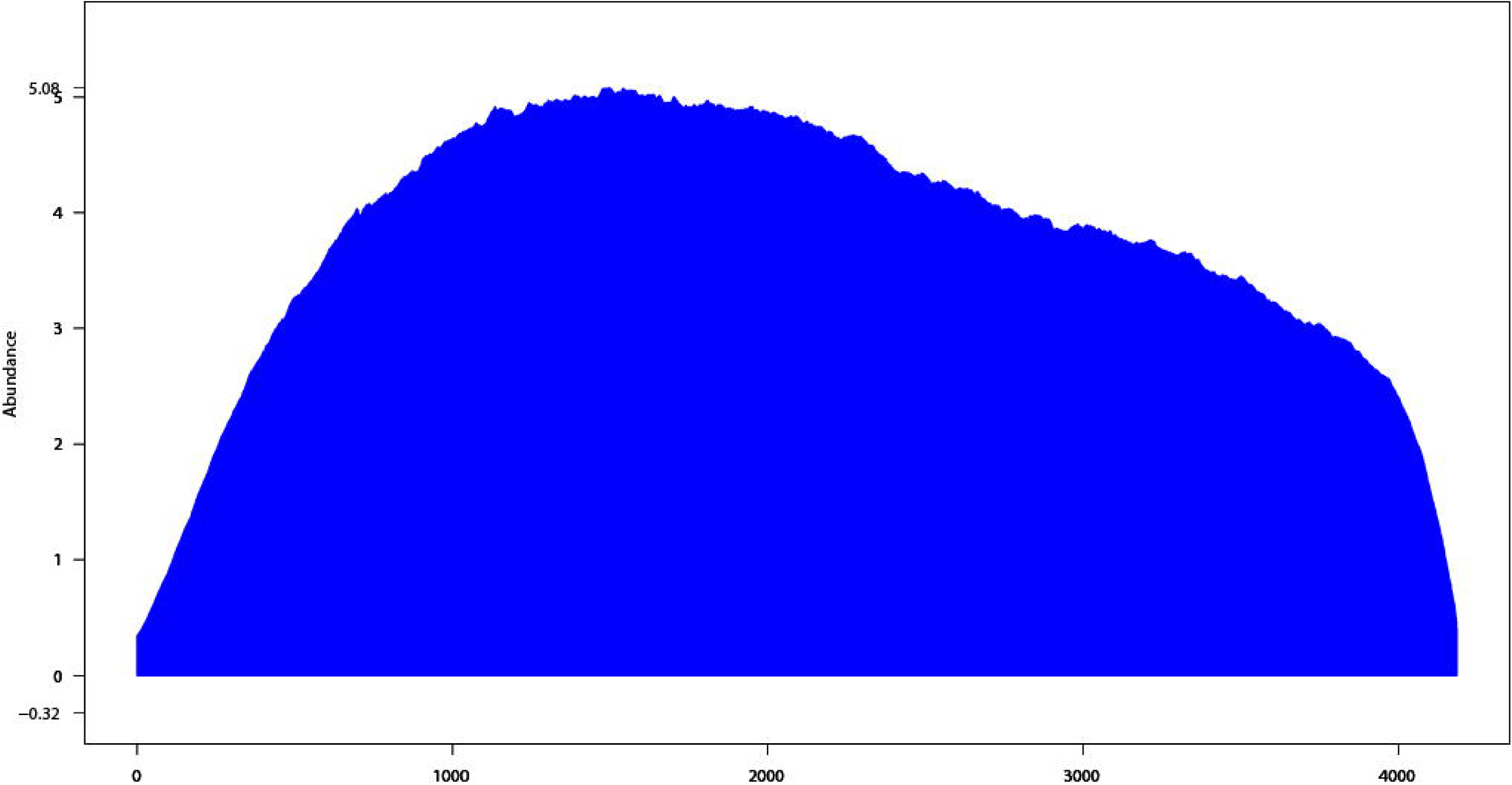
Metagene plots of RNA-seq at 5’ leader, CDS, and 3’ trailer of mRNAs. The x-axis shows the median length of all chicken mRNAs, and the y-axis represents the ppm (parts per million).

## DISCUSSION

The RNase H method is robust and capable of depleting rRNAs from low input samples [Adiconis, et al. (2013)]. Compared to Ribo-Zero, this method has a significantly higher efficiency in rRNA removal and slightly better evenness of coverage [Adiconis, et al. (2013)] as shown by the metagene analysis of a chicken RNA-seq results (Figure 3).

The whole procedure takes about 2.5 hours starting with purified RNA, and 3.5 hours starting with organic tissue. The procedure time is comparable to those of traditional methods, which take approximately 2 hours. The price of the method of interest would vary from $20 per reaction starting with total RNA (as seen in Table V) to $29 starting with organic tissues (seen in Table VI), while the cost of traditional methods varies from $55 to $118. By comparing the price and time with other traditional mRNA purification methods, as seen in Table VI, our method is cost-effective and efficient, while maintaining an excellent quality of purification.

**Table V:** Cost Estimation of the method of interest

| Name of Material/<br>Equipment | Quantity | Company | Catalog<br>Number | Comments/Description |
| --- | --- | --- | --- | --- |
| DNA Oligo | 32 mL | Integrated DNA<br>Technologies | N/A | \$912, \$5 per<br>reaction |
| TURBO DNase(2 U/uL) | 5000<br>units | Thermo Fisher | AM2239 | \$309, \$0.24 per<br>reaction |
| RNase H (2 U/μL) | 120<br>units | Thermo Fisher<br>(Invitrogen) | 18021071 | \$489, \$8.2 per<br>reaction |
| RNA Clean &<br>Concentrator Kits | 50 prep | ZYMO Research | R1013 | \$170, \$3.4 per<br>reaction |
| Acid Phenol Chloroform | 400 mL | Thermo Fisher | AM9722 | \$197, \$0.14 per<br>reaction |
| Glycogen | 1 mL | Sigma Aldrich | 10901393<br>001 | \$151, \$3.02 per<br>reaction |
| Total Cost Estimation | | | | \$2,228 Total<br>\$20 per reaction |

**Table VI:**
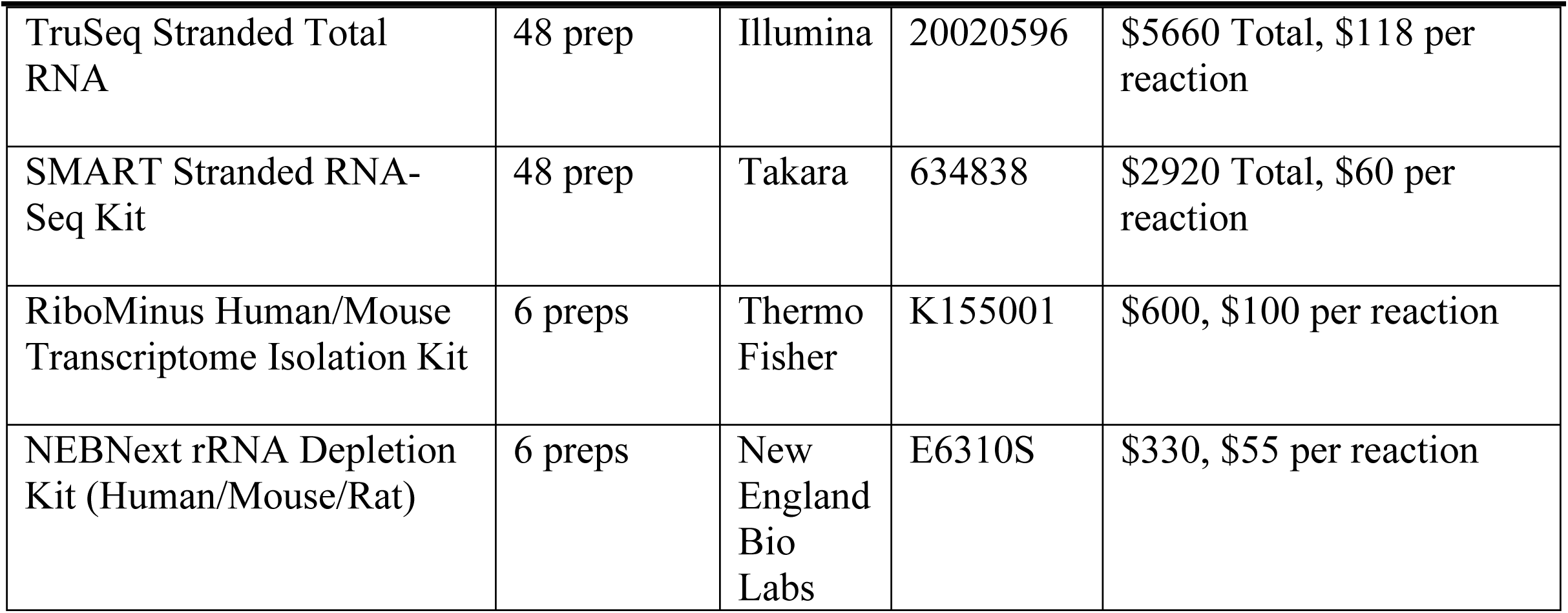
Other Methods cost Estimation

Depending on how the oligos are designed and which type of sequence the oligos are specified to target, the same method can potentially be applied for a wide range of organisms and different types of RNA. As long as the oligo is designed to antisense to the specific type of RNA of a specific species, this method should guarantee good quality of purification.

## ACKNOWLEDGMENTS

We thank N. Anthony for providing chicken tissues, UR Genomics Research Center for performing bioanalyzer analysis, and members of the Li laboratory for discussion. This work was supported by Agriculture and Food Research Initiative Competitive Grant no. 2018-67015-27615 from the USDA National Institute of Food and Agriculture to X.Z.L.

## Supplementary Files

**Supplementary Table 1:**
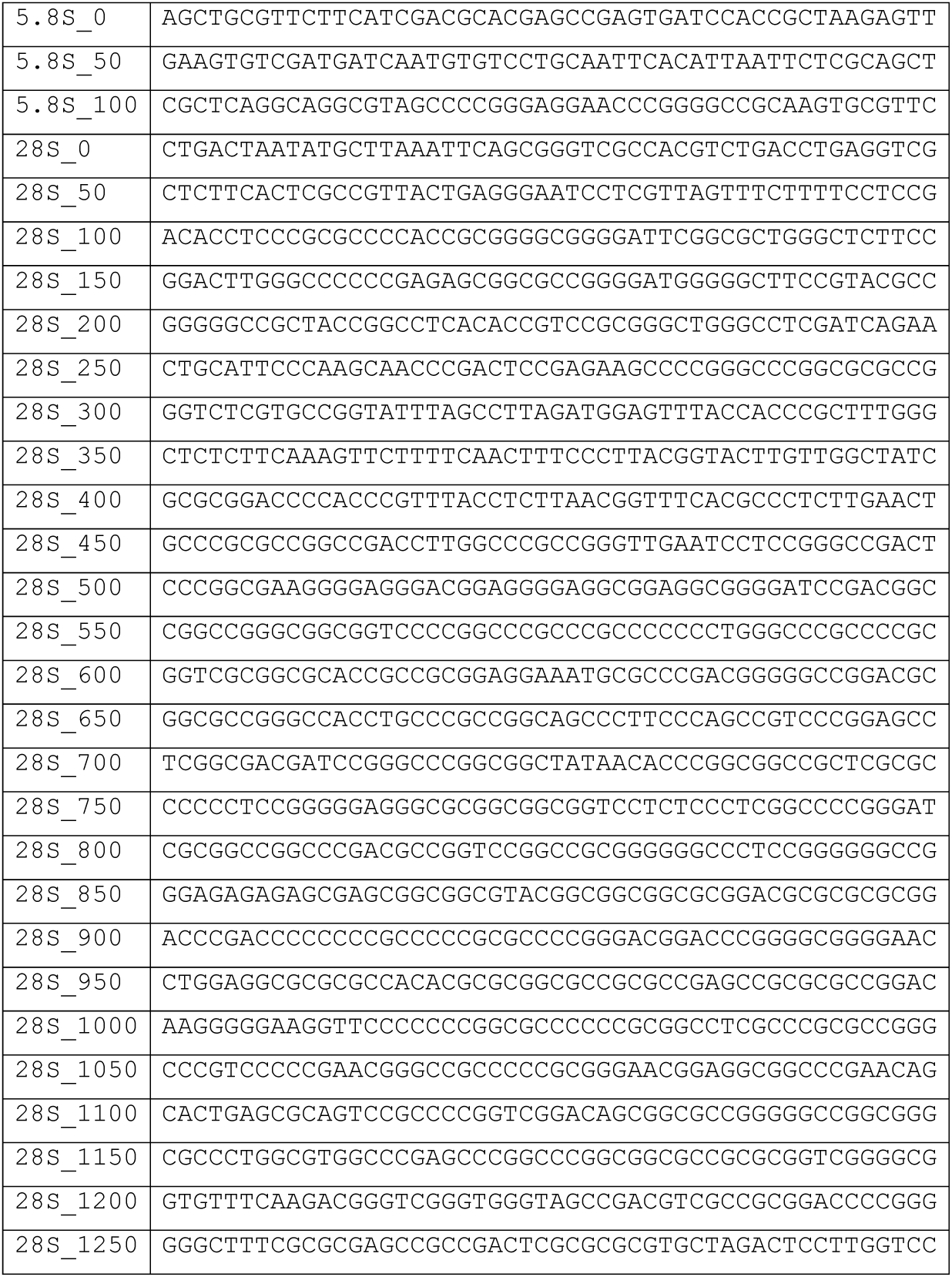
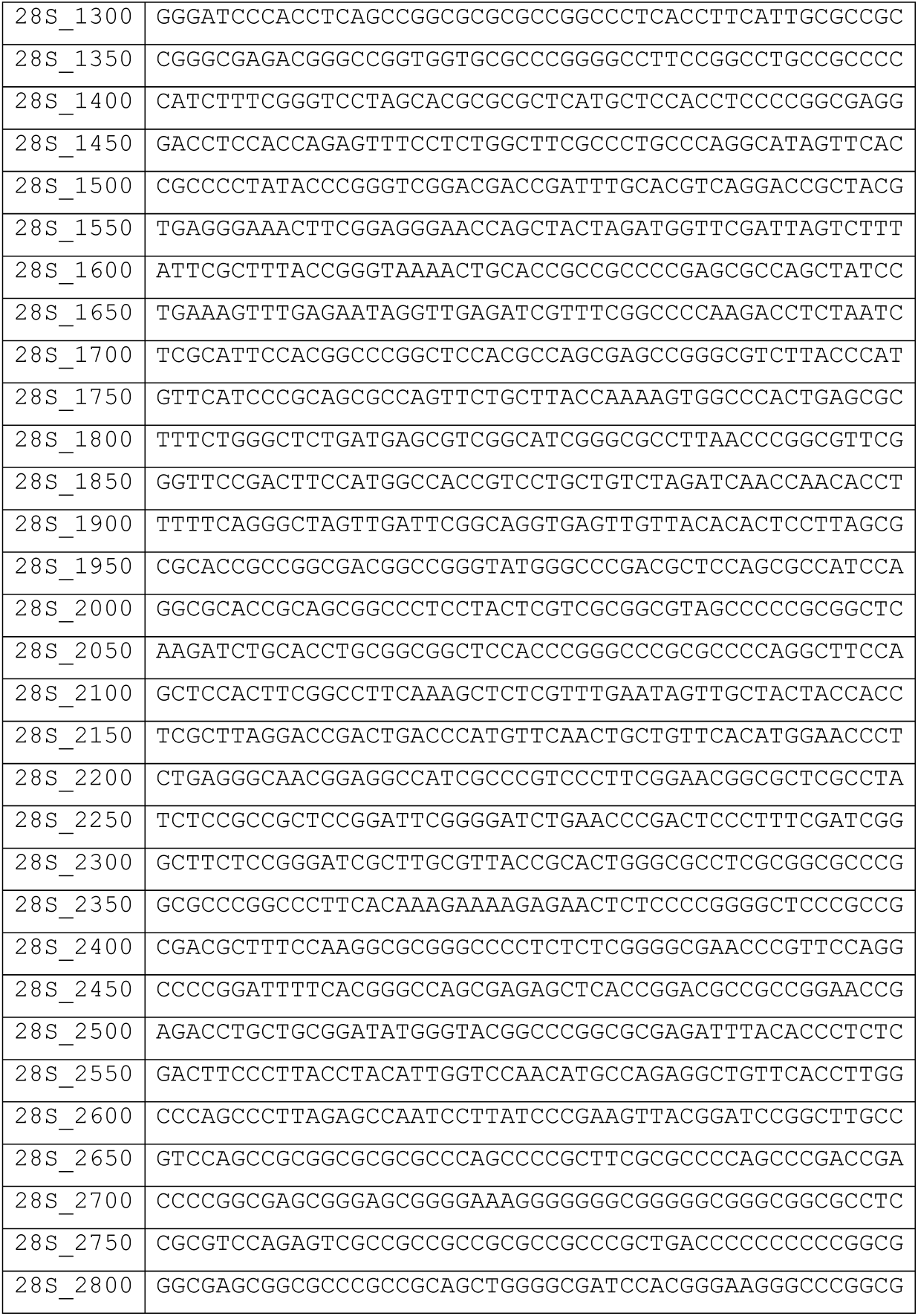
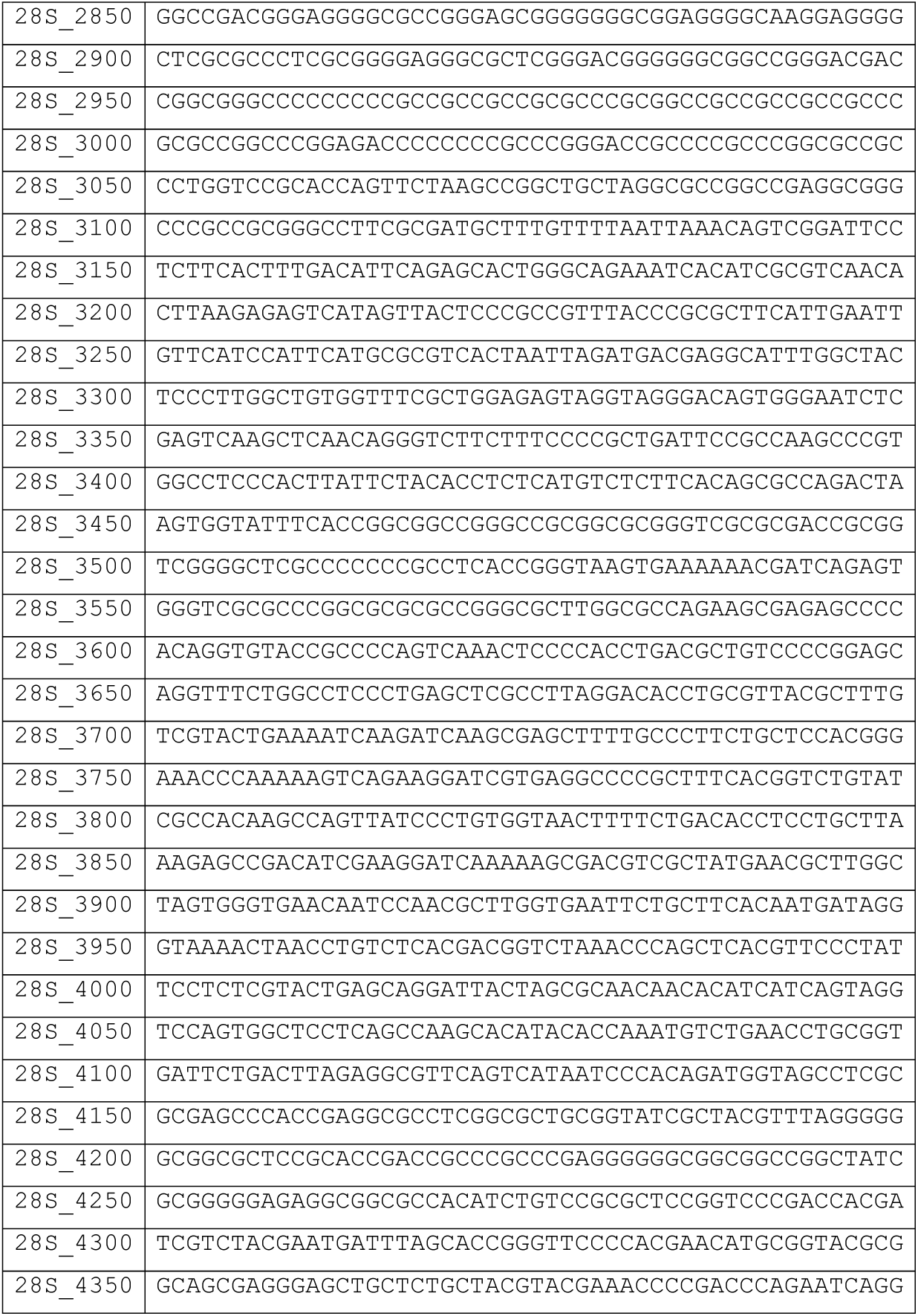
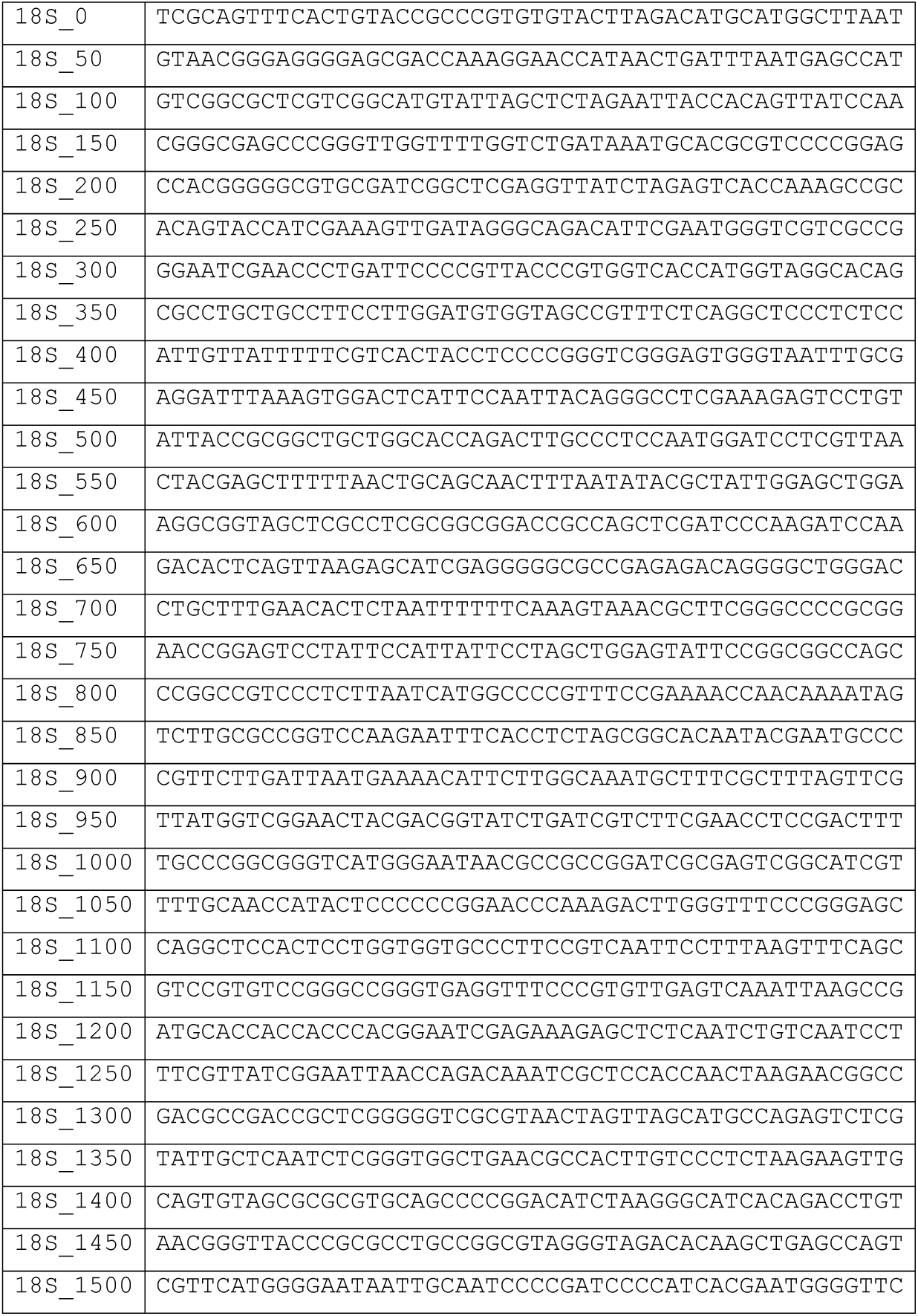
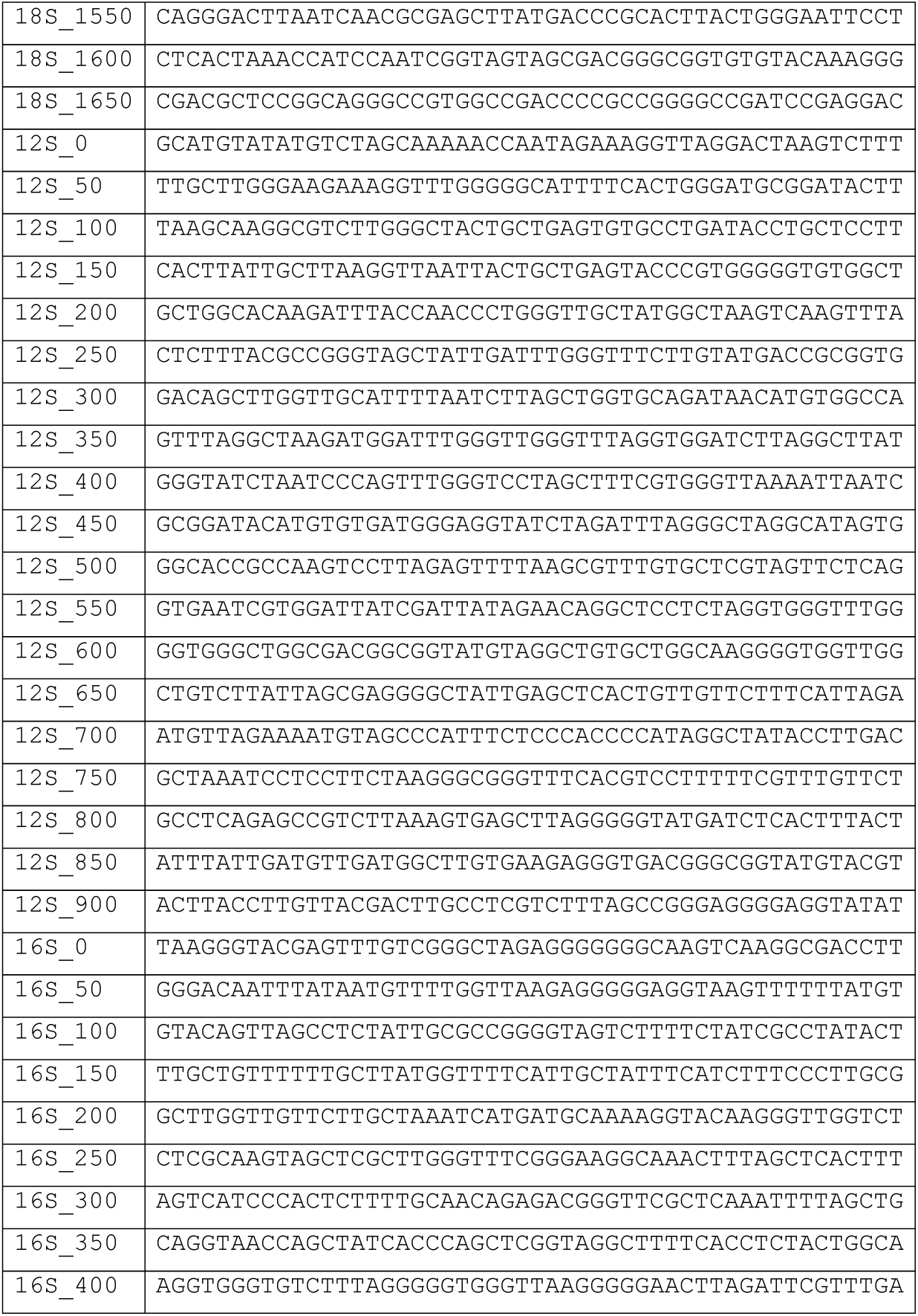
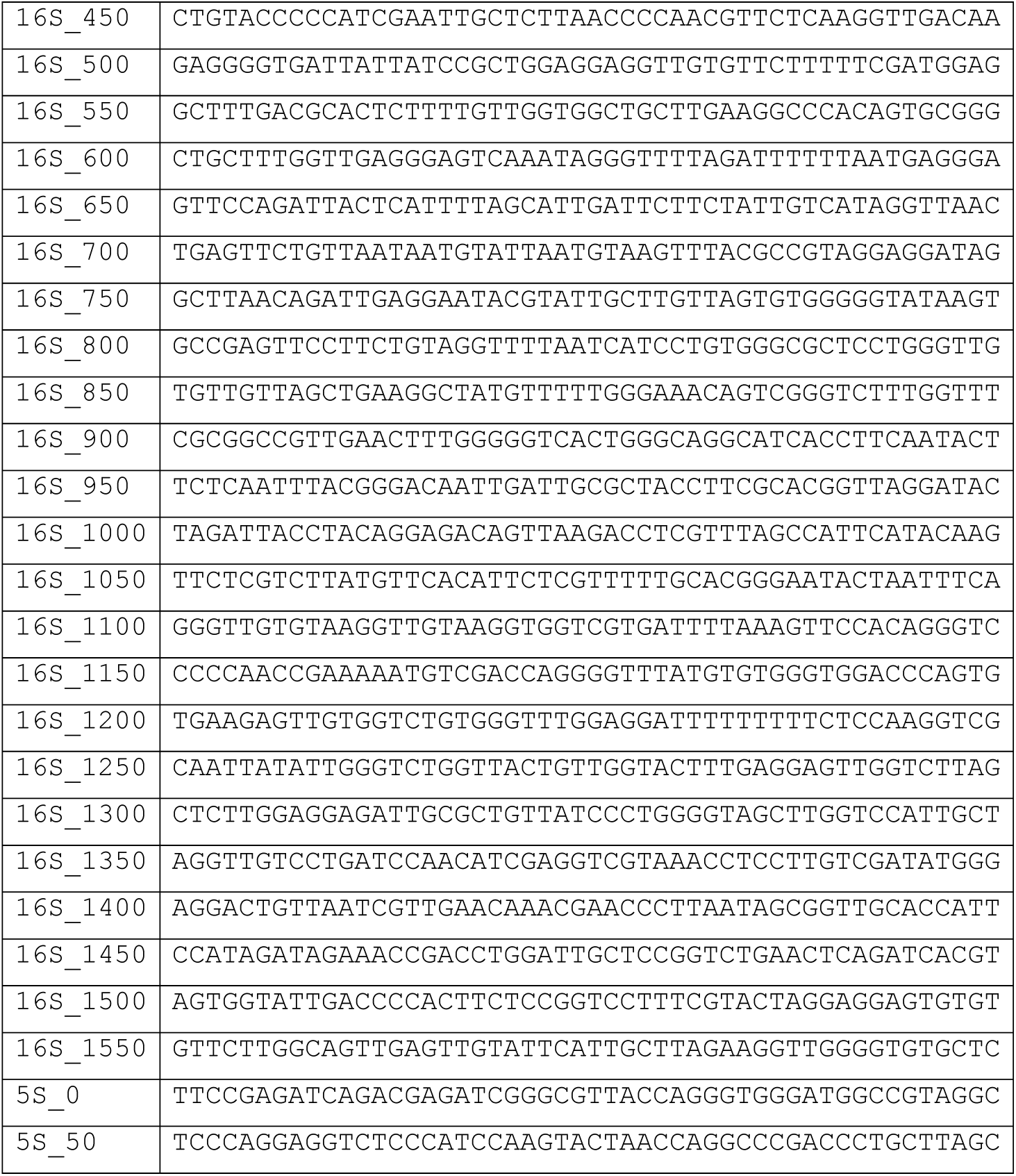
Sequence used for constructing Probe

